# Global urban forest cover and nature access is insufficient for human wellbeing and worsening

**DOI:** 10.64898/2026.09.10.750750

**Authors:** Robert I. McDonald, Mariami Marsagishvili, Lindsi Seegmiller, Peter Olsson

**Author notes:** Corresponding author: Correspondence to: Robert. I. McDonald.

## Abstract

Target 12 of the Global Biodiversity Framework urges governments to enhance human access to nature to improve human wellbeing. Here, we study nature access using several metrics for a globally representative sample of 140 large Functional Urban Areas, examining variation among regions and over time. We find most urbanites globally do not reach common benchmarks for urban forest cover and nature access. Globally, 3.3 billion people (76% of urbanites) live in urban areas that do not have at least 30% tree canopy cover. Moreover, 1.1 billion people (23% of urbanites) are farther than 1 km from a park as identified in Open Street Map, and 1.5 billion people (36% of all urbanites) are more than 1 km from a green patch as defined by land cover. Over time, urban nature access has declined. Between 1992 and 2020, most urban residents had an increase in the distance to green patches (63% of urbanites increased, average increase 680 m). Similarly, between 1992 and 2020, 31% of urbanites had a decrease in percent natural cover, while only 1.3% had an increase. We show that the variation among cities or regions in the amount of nature access depends on the metric used and the context. On average, mesic climates have more tree cover than do arid climates, and cities with lower GDP or higher population density tend to have lower natural share than other cities. Our results suggest that global urban nature access is insufficient for supporting human health and worsening over time.

## Introduction

Urban residents benefit from several ecosystem services that improve human well-being [1–3]. Exposure to urban nature has benefits to health, for instance, which occur through several routes [4–6]. One route operates through physical health, where parks encourage physical activity and recreation [7], which can help reduce obesity [8, 9]. A second route operates through mental health [10], where exposure to nature has been shown to reduce stress [e.g., 11, 12, 13], improve self-reported mental health [14], and reduce the incidence of depression [15] and schizophrenia [16].

There are multiple types of nature in cities [17], that each support human well-being by providing different ecosystem services in different amounts [18, 19]. This study focuses on two main types of urban nature, urban tree canopy and urban green spaces such as parks. Urban tree canopy provides a number of benefits, including cooling the air through evaporative cooling [20, 21], providing aesthetic beauty [22], and improving stormwater drainage [23]. Parks and other open space also provides a number of benefits, most notably recreational and physical health benefits [19] and harboring biodiversity [24].

Perhaps because of the vital role that urban forests and green space (which we will collectively refer to as “urban nature”) play in maintaining human well-being, many governments are trying to increase urban nature access and exposure for their citizens. At a global level, this is reflected in the Convention of Biological Diversity Kunming-Montreal Global Biodiversity Framework (GBF), in which national governments around the world commit to protect and restore biodiversity and natural systems [25]. Specifically, Target 12 of the GBF commits governments to “significantly increase the area and quality and connectivity of, access to, and benefits from green and blue spaces in urban and densely populated areas sustainably… thereby improving human health and well-being and connection to nature…” [26]. Since the target was written, there have been guidance and suggested methodologies provided for how to measure progress toward Target 12 by working groups under the CBD [26].

In scientific literature, there have been several studies that have gone beyond the general directional goal of Target 12 (i.e., increasing urban nature area and access), but have suggested specific thresholds of how much nature is needed to maintain well-being. For urban forest cover a threshold of 30% has been suggested as adequate to maintain well-being [27, 28] and biodiversity [29]. This 30% thresholds comes in part from the empirical dose-response modeling by Cox and colleagues [27], which found 50% less depression and 43% less stress in neighborhoods with more than 20% forest cover and 56% less anxiety in neighborhoods with more than 30% forest cover. Another commonly suggested threshold is that all residents should be within a short walk of a park or other green space (15min, or sometimes 1km). This threshold is related conceptually to the goal of the 15-minute city (Moreno et al. 2021), which is considered the length of time a short walk from one’s house would take. Multiple variables are sometimes combined into guidelines for planners, such as the 3/30/300 guidelines [30], which suggest urban residents should be able to see 3 trees from their dwelling, live in a neighborhood with 30% tree cover, and live within 300m of a green space. The effect of hitting this threshold has been evaluated for its empirical effects, such as on mental health [31], and recent studies have quantified how well some cities are doing toward achieving this rule [32].

Despite this growing literature on how much nature is needed to ensure well-being, it remains unclear how much access or exposure the average global urban resident has relative to the suggested thresholds needed for human well-being. In part, this is because most multi-city studies have looked at a biased sample of urban areas globally [33], such as by only examining cities in the United States [34] or Europe [35], or preferentially focusing on large, mega-cities [36]. For instance, Bertassello et al. [32] recently assessed a large sample of 862 European cities against the 3-30-300 rule. In part, it is because most global multi-cities studies have focused on single metrics, such as the study by Wei et al. [37] of the area of urban green space relative to a city’s population. We address both gaps in this study by calculating and comparing multiple metrics of urban nature for a globally representative sample of the world’s urban areas.

Trends over time have also been difficult to assess, leaving us unsure of how urban nature access is changing for residents. The ultimate goal of Target 12 is to achieve change in urban nature access over time, but for some measures of urban nature it can be challenging to assemble a time series of consistent data [38]. While there have been some studies of global urban nature exposure over time, most notably using tree cover [39] or indices of greenness [40], there is relatively little understanding of global trends in urban access to parks and green space over time, either geographically or temporally. We address this gap in this paper by assessing both urban tree cover and access to parks and green space over time, for a globally representative sample of cities.

Globally assessing urban nature access is challenging because there are several contextual variables that determine current levels of nature access and future plausible targets. For instance, climate, particularly aridity, controls whether trees can grow without supplemental watering, and so climatic gradients correlate strongly with urban tree canopy cover [19, 34, 41]. Additionally, more densely settled neighborhoods, with a higher population density and impervious surface cover, tend to have less tree cover. Higher population density and other aspects of urban form are in turn products of historical processes and the city’s economic status— urban areas in developing economies are typically denser than in developed economies [42]. To address the differing contextual variables of a global study, we measure climate gradients and urban density alongside our different nature access metrics.

In this paper, we aim to partially close these research gaps discussed above by taking a statistically representative sample of 140 Functional Urban Areas, stratified across geographic regions, and then assemble multiple datasets that measure urban nature access in diverse ways and at various times. We then assessed the following research questions:

1. How is urban access to nature compared to proposed benchmarks for safeguarding human health and well-being?
2. What is the current trend over time in urban residents’ access to urban nature?
3. How much does the metric used to assess urban nature access influence the global patterns observed?
4. How do contextual factors such as climate and level of economic development correlate with residents’ access to urban nature?

## Materials and Methods

Our methods proceeded in four main steps. First, we chose an urban area definition and selected a representative sample of urban areas. Second, we assembled geospatial data for multiple time points, where available, for that sample of urban areas. Third, we conducted a GIS analysis to measure urban nature access in multiple ways, as well as various proxy measures of urban form and context. Fourth, we conducted descriptive and statistical analyses using these variables to answer the research questions listed in the Introduction. Below, we discuss each of these four stages on our methodology in more detail.

### Urban areas studied

We follow the Functional Urban Area (FUA) definition [43]. This definition was applied in a consistent manner globally and includes both the central city and contiguous commuting zones. Urban area polygons were obtained from the Global Human Settlements Layer’s 2019 boundary file (R2019A)[44], which was estimated using 2015 population data. Note that the GHS Urban Centre Database 2025 [45], based upon 2023 population data, was released as this manuscript was being prepared, and so was not available for use in our GIS analysis. As the urban population has grown between 2015 and 2023, the latter urban boundary is slightly wider, but similar in shape and form, and so we believe the distinction between the two FUA definitions would be minor for the results we present in this manuscript.

The goal of this study was to consistently estimate average nature access, using multiple metrics, for large urban areas globally and by regions. Our definition of regions was the World Bank’s 7 regions: East Asia and Pacific, Europe and Central Asia, Latin America and the Caribbean, Middle East and North Africa, North America, South Asia, Sub-Saharan Africa [46]. Large urban areas were defined as those having a FUA population (core plus commuting area) greater than 1 million. We chose to focus on large urban areas due to data quality limitations, since Open Street Map data were found in preliminary examinations to be more complete and spatially detailed in large urban areas. Finally, we randomly selected a sample of 20 large FUAs from each of the 7 regions (N=140). We stratified across regions to have a representative sample of FUAs for each region, so we could study differences among regions. The number of 20 FUAs per region was chosen to have enough replication to enable consistent estimation of average nature access, by region and globally, for a variety of metrics, based upon preliminary analyses of the stability of the estimated mean for contextual variables (see below) for FUA samples drawn from different sizes.

### Data Sources

#### Landcover

The finest resolution landcover data used was the WorldCover 10m dataset [47]. This dataset has the advantage of being one of the highest resolution global land cover datasets that are freely available globally. The disadvantage is the short duration for which the WorldCover data is available, starting in 2020. For this study we use just one timepoint for this data source, from 2021, for which the WorldCover v200 dataset [47] was retrieved from Google Earth Engine. The WorldCover files were processed in R version 4.5.3 to derive percent coverage of both tree and natural land cover classes within each FUA boundary. A single land cover class code, 10, was used to define tree land cover. The following land cover class codes were defined as natural cover: 10, 20, 30, 90, 95, 100.

In order to analyze historical changes in access to nature, the ESA-CCI dataset was used [48]. It has a relatively coarse resolution of 300m but is available annually from 1992 to 2020. In this study, we use the first and last year of this series. As with the 10m land cover, we looked at tree cover and natural land cover but subdivided natural land cover into blue and green natural land cover types. Tree cover was defined to include the following ESA-CCI land cover class codes: 40, 50, 60, 61, 62, 70, 71, 72, 80, 90, 160, 170. Green cover was defined as including the following land cover class codes: 40, 50, 61, 62, 70, 71, 72, 80, 90, 100, 110, 120, 122, 130, 150, 152, 153, 160, 170, 180. Blue cover was defined to include only permanent water bodies, class code 210.

#### Open Street Map parks

To extract information on land use and protection, we used the osmdata library [49] in R to extract OpenStreetMap (OSM) data for parks and green spaces. While these data are correlated with land cover, there are crucial differences. Public parks may not always be represented in land cover datasets as natural land cover codes (e.g., playgrounds might contain a mix of impervious, sparse vegetation, and tree cover), while privately owned green spaces would be captured in land cover datasets but not in the OSM data. We chose to extract parks and other similar areas from OSM, rather than using the World Database on Protected Areas (WDPA) or similar datasets of strictly protected areas, since preliminary investigations in many urban areas showed that many urban parks and green spaces are not listed in the WDPA, particularly those that are municipally owned or managed. OSM contains many distinct kinds of features with tags (or keywords) that reflect their use. To extract park features of interest we developed and used a set of OSM tags commonly used in our 140 sample FUAs. Within the ‘Leisure’ attribute of the OSM dataset, we extracted features with the tags: *park, nature_reserve.* Within the ‘Natural’ attribute, we extracted features with the tags: *valley, beach, hill, dune, cliff, wetland, coastline, bay, wood, treerow, shrubbery, grassland, forest.* Within the ‘Landuse’ attribute, we extracted features with the tags: *forest, meadow, grass, park, recreation_ground, nature_reserve, wood.* Within the ‘Boundary’ attribute, we extracted features with the tag *protected_area.* If a feature contained any of these tags, we included it in our analysis.

#### Other contextual variables

##### Population

We used WorldPop [50], which has globally available population estimates at 1km resolution, beginning in 2000 and continuing every five years until the present. We chose the WorldPop 1km population data over other datasets because it has consistently estimated grids of population every five years over the time span of interest for this project.

##### Per-capita GDP

Our estimate of economic development, as measured in per-capita GDP, was taken from Kummu et al. [51], which assembled estimates of per-capita GDP (in USD PPP) at sub-national scales between 1990-2022 and then multiplied by population to provide grid datasets of GDP over time. In this paper, we are using the estimates of per-capita GDP, and for each FUA we calculated the population-weighted mean per-capita GDP.

##### Aridity Index

To measure how dry or mesic a climate is, we used the Aridity Index (AI), as defined by the UN Environment Programme (UNEP) as the ratio of precipitation to potential evapotranspiration [52]. Values below 0.20 are considered arid, values between 0.2-0.5 are considered semi-arid, values between 0.5-0.65 are dry subhumid, and values above 0.65 are considered humid climates. Specifically, we used the 30 arcsec resolution (v3) maps of AI [53].

### Analysis

#### GIS Analyses

We created a common 1km grid across all 140 FUAs using R and the *sf* (spatial features) package [54] using the Eckert IV equal-area projection. This grid served as the spatial framework for our analysis. For each 1km grid cell, we calculated the mean values within the cell for all natural access variables described below.

To analyze the distance to natural patches for the years 1992 and 2020, the ESA-CCI 300m data were reclassified into categorical rasters, distinguishing between blue zones and green zones, and the Euclidean distance calculated from all cells to the nearest zones. In a similar fashion, the Euclidean distance to OSM park spaces was calculated. In a few FUAs, there was no water body within the FUA, so distance to water was undefined for those FUAs. In these few cases, we recorded a value of distance to blue that represents the maximum Euclidean distance from the center of the FUA to the edge of each urban area. We chose to do this because the alternative (treating this variable as undefined in these few cases and excluding them from the analysis) would potentially introduce a bias, since these few cases that have a population that is very far from water would be excluded from our analysis.

### Descriptive and statistical analysis

To calculate average summary statistics at the FUA level, we used the population-weighted average. This is important since different pixels in a FUA have vastly different populations and we wanted to measure the average individual’s access. The urban core is relatively small, having more people and relatively fewer natural features than the commuting zone, which is bigger but has fewer residents and relatively more natural features. The population-weighted average allows us to estimate the average amount of urban nature access experienced near people’s homes, rather than having the average dominated by the value in pixels in the commuting zone. When aggregating up to regional average, we used the simple average of the FUAs in each region to estimate the mean and standard error of the estimate. When aggregating up to global estimates, we used the regional estimates, but each region was weighted by its total urban population, as estimated by the UN Population Division. This accounts for the fact that there are more urban residents in the East Asia and Pacific region, for instance, than in the Latin American and Caribbean region. Our estimate of the standard error of global means accounts for the uncertainties in each regional estimate.

To evaluate correlations between different variables, we used Pearson correlations, while for hypothesis testing between variables, where we are hypothesizing that one variable causally relates to another, we used simple linear regression. Where relevant, the assumption of a linear relationship was visually checked in a scatterplot first, although with our relatively small sample size (n=140) the data, while sometimes noisy, did not depart visually from a roughly linear relationship. For this reason, we did not transform any of the variables used in our analysis, instead using the original variables.

## Results

### Urban nature access

Urban nature access varies widely across urban areas. For instance, using the metric of the distance to the nearest green patch calculated from the ESA-CCI 300m data, the population-weighted average distance varies from less than 500m to more than 3,000m (Figure 1). In general, FUAs in North America, Europe, and South America have shorter average distances to nature (i.e., more nature access), while FUAs in Africa, the Middle East, and Asia have longer average distances. Within Asia, FUAs in the Indian subcontinent have greater average distances than FUAs in eastern Asia.

**Figure 1.**
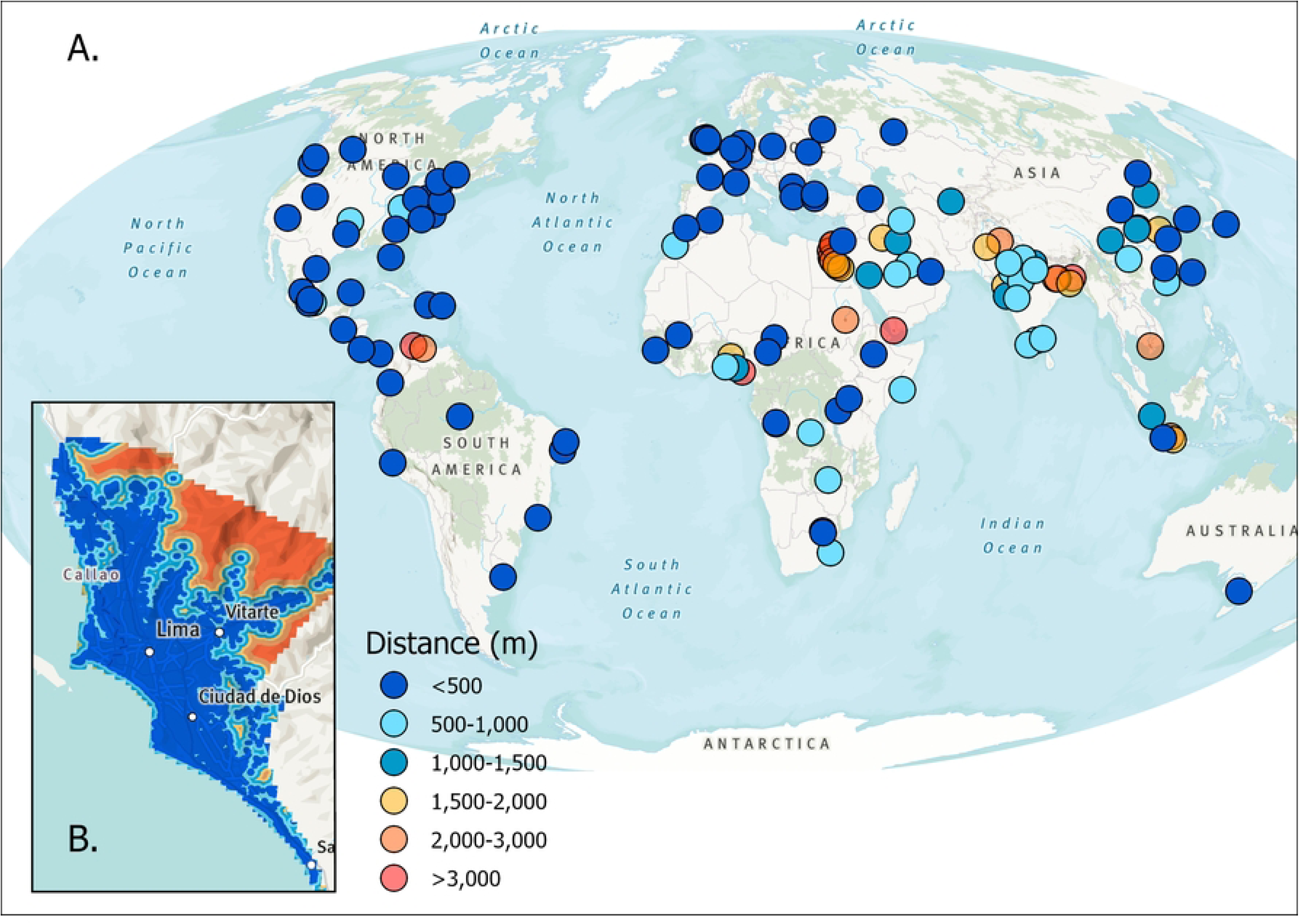
Distance to a green patch. A.) Average population-weighted distance to green patch for a globally representative sample of 140 Functional Urban Areas (FUA). B.) A close-up of the Lima FUA, to give a sense of the underlying data used to calculate the population-weighted average. Note that the legend is the same between the two panels, with blue indicating shorter distances and red indicating greater distances.

Similar global and regional patterns are observed with other metrics of nature access (Table 1). Measured by natural cover or tree cover, North America exhibits the greatest nature access (at 43% and 32% respectively), followed by the Europe and Central Asia region (at 38% and 26% respectively). Conversely, the Middle East and North Africa region have the lowest natural cover (10%) and tree cover (5%). Patterns for distance to parks, as defined in the OSM, are slightly different than for natural land cover and tree cover (Table 1). The Europe and Central Asia region is the region with the shortest distance to OSM parks (distance of 180m), followed by North America (260m). Like patterns with nature cover and tree cover, the Middle East and North Africa region have the longest distance to OSM parks (2,000m).

**Table 1.** Current population-weighted access to nature, by World Bank region. Shown is the natural land cover (%), the tree cover (%) and the average distance to parks in Open Street Map (OSM) as of 2025. Land cover is derived from 10m WorldCover data (2021).

| WB Region | Natural Cover (%) | Tree Cover (%) | Distance to OSM Parks (m) |
| --- | --- | --- | --- |
| East Asia and Pacific | 27.5 | 23.3 | 1,011 |
| Europe and Central Asia | 38.1 | 26.4 | 184 |
| Latin America and Caribbean | 23.2 | 14.9 | 514 |
| Middle East and North Africa | 10.3 | 5.1 | 2,009 |
| North America | 43.3 | 31.7 | 260 |
| South Asia | 26.2 | 20.3 | 1,443 |
| Sub-Saharan Africa | 16.4 | 9.2 | 880 |

Existing urban tree cover or natural cover are generally insufficient to meet suggested benchmarks for most urbanites (Table 2). For instance, 30% is a common benchmark for adequate urban tree cover for mental health and wellbeing [27, 28, 55], yet we estimate that globally, 3.3 billion people (76% of all urbanites) live in urban areas that do not meet this target. If we assess instead a 30% benchmark for natural cover, we estimate that globally 2.8 billion people (64% of all urbanites) live in urban areas that are below this benchmark.

**Table 2.** Global average urban access to nature and the proportion of people below a benchmark. To estimate global urban access, we accounted for each region’s total urban population when calculating the average indicator value and the proportion of people below a benchmark.

| Metric | Global average |  | Benchmark | Proportion of urbanites failing to meet benchmark |  |
| --- | --- | --- | --- | --- | --- |
|  | *Mean* | *Standard error* |  | *Mean* | *Standard error* |
| Tree cover | 19% | 3% | >30% | 76% | 7% |
| Natural cover | 27% | 3% | >30% | 64% | 11% |
| Distance to OSM park | 905m | 199m | <1000m | 23% | 8% |
| Distance to green patch | 1,910m | 525m | <1000m | 36% | 9% |
| Distance to blue patch | 5,105m | 503m | <1000m | 98% | 1% |

In contrast, most urbanites meet suggested benchmarks for park or green space access. We assessed the proportion of people that are within 1km of an OSM park, which roughly corresponds to the area within a 15-minute walk of a park, a common goal for park access [56]. We estimate that globally 1.0 billion people (23% of all urbanites) are farther than 1km from an OSM park. If we use distance to green patches, as defined from the 300m resolution land cover, the fraction of urban dwellers failing to meet the 1km benchmark is similar but slightly greater (36% of all urbanites) The situation is quite different for access to blue spaces, with the vast majority (98%) of urbanites more than 1km from a blue patch.

### Trends over time

Trends over time in urban nature access can only be assessed with 300m resolution land-cover derived metrics since it is not generally possible to analyze change over time with the OSM dataset. At this scale, only large patches of natural habitat can be observed, while small urban parks present in the OSM dataset are not observed in the dataset. Distance to large green patches generally increases over time (Table 3), with the South Asia region experiencing the largest increase, which is consistent with the more rapid urban growth in this region, which is converting natural habitat to other developed uses.

**Table 3.** Distance to green and blue patches over time, by World Bank region. Shown is the distance to blue and green patches (derived from 300m data ESA-CCI data) for the years 1992 and 2020.

| WB Region | Distance to Blue Patches (m) |  | Distance to Green Patches (m) |  |
| --- | --- | --- | --- | --- |
|  | 1992 | 2020 | 1992 | 2020 |
| East Asia and Pacific | 7,118 | 6,474 | 1,602 | 1,898 |
| Europe and Central Asia | 4,674 | 4,030 | 944 | 984 |
| Latin America and<br>Caribbean | 9,207 | 6,694 | 655 | 845 |
| Middle East and North<br>Africa | 8,206 | 5,121 | 1,109 | 2,346 |
| North America | 5,457 | 3,943 | 510 | 552 |
| South Asia | 4,457 | 3,356 | 2,079 | 4,903 |
| Sub-Saharan Africa | 7,210 | 4,024 | 599 | 748 |

In 2020, average distance to green patches (i.e., terrestrial natural land covers) is smallest in North America (550m distance to nearest green patch on average), followed by Sub-Saharan Africa (750m) and Latin American and Caribbean (850m). South Asia has the farthest average distance to green patches (4,900m).

Distances to blue patches show a general decreasing trend. One contributing factor is that a few features were classified as blue in 2020 but not in 1992. The Latin America and Caribbean region shows the longest distance to blue patches in both years, with a reduction from 9km in 1992 to 7km in 2020. This trend is mirrored in the Middle East and North Africa, where blue distances decreased from 8km to 5km. Conversely, East Asia and Pacific as well as Europe and Central Asia show no dramatic change through these years, with an average distance changing from 7km to 6km and 5 to 4 km.

In 2020, people are nearest to blue patches in the South Asia region (3,400m average distance to blue patch), followed by the North American region (4,000m). People in the Latin America and Caribbean (6,700m) and East Asia and Pacific (6,500m) regions are furthest from blue patches on average.

Over the period studied, most urban areas have increased in population density, and urban residents are farther from large green patches (i.e., have less nature access) (Figure 2). The North America (NA) and Europe and Central Asia (ECA) regions show the least change in population density and the smallest increase in distance to nature (i.e., the smallest decline in nature access). Conversely, South Asia (SA) and the Middle East and North Africa (MENA) regions show the largest increase in distance to green patches (i.e., the greatest decline in nature access), with South Asia increasing from 2.0km meters to 4.9km, and MENA increasing from 1.1km to 2.3km meters. Sub-Saharan Africa (SSA) shows the greatest increase in population density during these years, from 2,100 to 4,000 people per square kilometer, while the distance to nature has only slightly increased from 600m to 750m.

**Figure 2.**
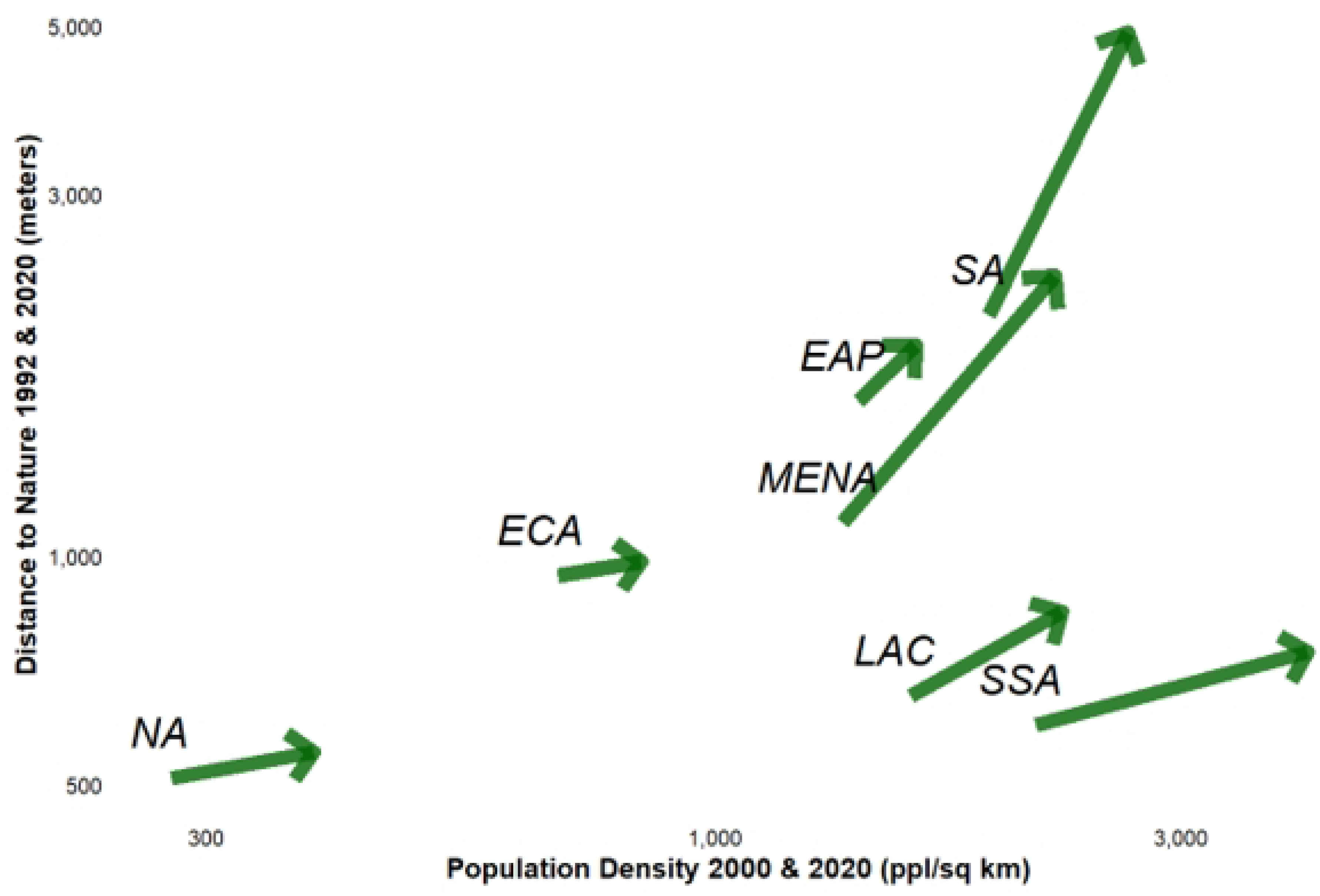
Change over time in population density (people per square km) and distance to nature based on ESA-CCI landcover (meters) from 2000 to 2020 across various World Bank regions. The following acronyms were used: North America: NA; Europe and Central Asia: ECA; East Asia and Pacific: EAP; Middle East and North Africa: MENA; Latin America & Caribbean: LAC; Sub-Saharan Africa: SSA; South Asia: SA. Green arrows indicate the direction of change over time for each region from 1992 to 2020.

Trends over time in land cover derived from the ESA-CCI data also show a general trend of decline over time (Table 4). Natural cover declines in every World Bank region, with the greatest declines in Sub-Saharan Africa and the Middle East and North Africa regions. Conversely, the smallest declines were in South Asia and the Europe and Central Asia region. Similarly, tree cover declines in every World Bank region, with the greatest declines in Sub-Saharan Africa and North America regions. Conversely, the smallest declines were in the Middle East and North Africa region and South Asia. Note that for arid regions, desert and arid land cover types were considered natural but do not contribute to the tree cover calculation. This explains why the Middle East and North Africa region can have relatively rapid declines in natural cover, but only a small decline in tree cover (i.e., there is relatively little tree cover in this region).

**Table 4.** Average population-weighted change natural cover and tree cover, derived from 300m data ESA-CCI data, between 1992 and 2020. Shown is the difference in percentage points (pp), with a negative number implying a decline in the land cover. Note that the coarser scale of ESA-CCI data leads to generally lower detection of natural and tree cover than the Worldcover data shown in Table 1, so the results are not directly comparable.

| **WB region** | **Δ Natural**<br>**Cover, 1992 to**<br>**2020 (pp)** | **Δ Tree Cover,**<br>**1992 to 2020**<br>**(pp)** |
| --- | --- | --- |
| East Asia and Pacific | -7.1 | -2.7 |
| Europe and Central Asia | -4.8 | -2.8 |
| Latin America and Caribbean | -16.5 | -3.9 |
| Middle East and North Africa | -20.5 | -0.7 |
| North America | -18.1 | -5.5 |
| South Asia | -3.7 | -1.4 |
| Sub-Saharan Africa | -29.1 | -9.5 |

### Nature access depends on the metric used

On the one hand, there are strong relationships (whether positive or negative) among many metrics of nature access and exposure (Figure 3). The information about relative nature access when comparing among FUAs is then similar for many metrics. On the other hand, the absolute value of two different metrics can be quite different. This may matter when comparing nature access to some fixed benchmark, or in terms of a policy goal of enabling universal adequate urban nature access.

**Figure 3.**
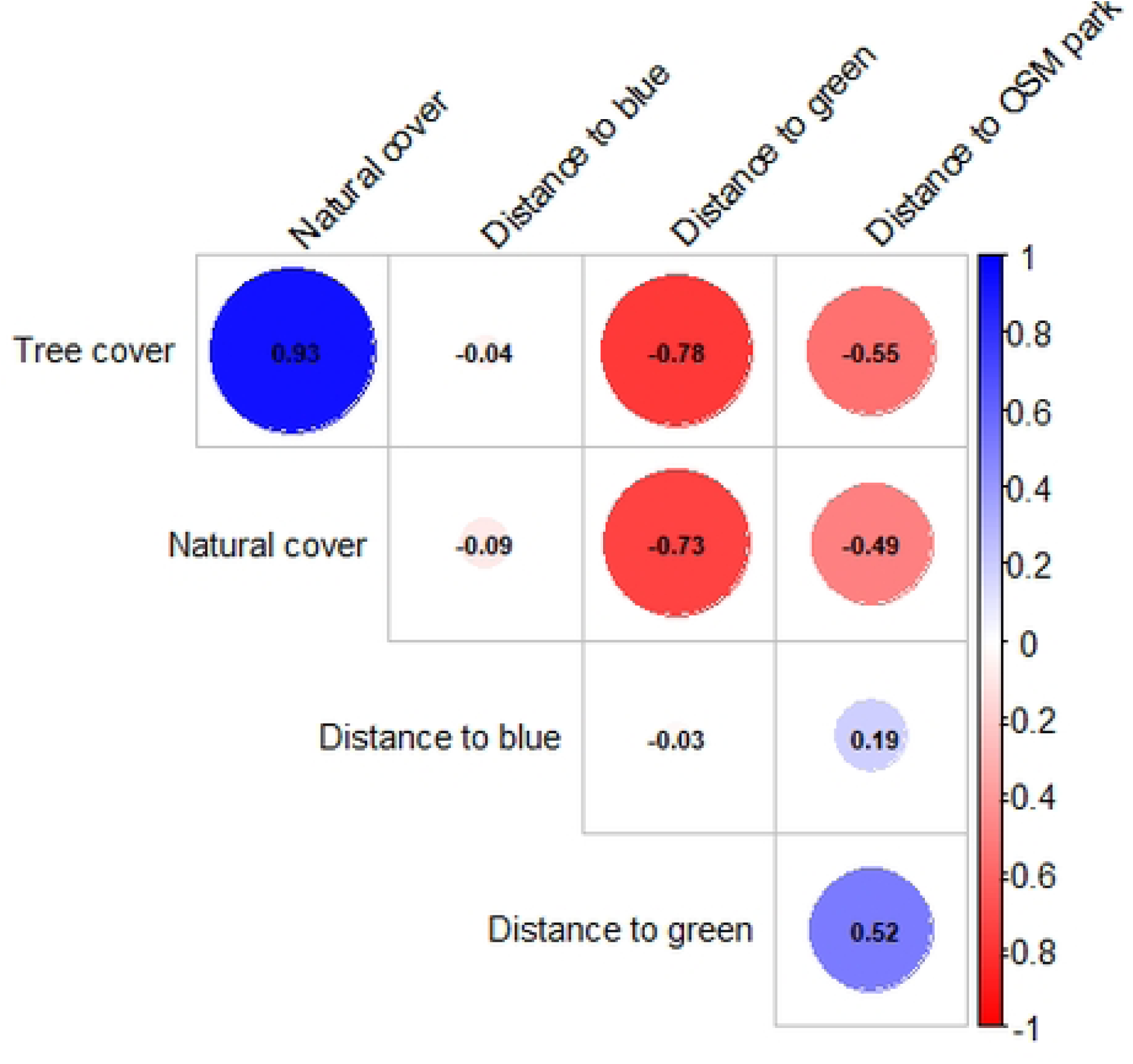
The Spearman rank correlation global nature access for the 140 cities in our sample, comparing global nature access metrics for 2020. Natural cover and tree cover are assessed with Worldcover data (2021), distance to blue and distance to green are assessed with ESA-CCI data (2020), and distance to OSM park is for available data in 2025.

For instance, the strongest rank correlation is between natural cover and tree cover (ρ= 0.93), which is expected since tree cover is one of the land cover categories included within natural cover. The biggest difference between natural cover and tree cover occurs in arid regions, such as the Middle East and North Africa, where the natural land cover type is predominately arid and semi-arid habitats that are not forested. Measured by natural cover or tree cover (Table 1), North America exhibits the greatest nature access (at 43% and 32% respectively), followed by the Europe and Central Asia region (at 38% and 26% respectively). Conversely, the Middle East and North Africa region has the lowest natural cover (10%) and tree cover (5%). Thus, while the two metrics are strongly correlated, the choice of using tree cover instead of natural cover means that in absolute terms, the amount of nature exposure will be substantially less.

Similarly, the metric of distance to OSM park is rank correlated with the distance to green patches (ρ= 0.52), as defined with 300m resolution land cover data. However, the rank correlation is less strong, possibly because there is a difference in the detection of small urban parks and parks without much natural land cover. In general, the distance to OSM park is less than the distance to green patches for most cities. The Europe region has a lot of small urban parks, and so nature access for this region looks relatively different for these two metrics.

Finally, it is notable that there are negative correlations between metrics of cover (tree or natural) and metrics of distance (green patch or OSM park), with rank correlations between ρ=-0.49 and ρ =-0.78. This is to be expected, since landscapes with a greater natural fraction will tend to have more and larger natural patches, and other portions of the landscape will tend to be closer to a natural patch.

### Urban context and urban nature access

The climatic context of a city affects tree cover. There is a strong positive relationship between the Aridity Index and tree cover (R=0.62, p < 0.001). An increase of 0.1 units of the Aridity Index (i.e., a wetter climate) correlates with a 3.5% increase in tree cover on average. There are large regional differences in aridity which correlate with differences in tree cover, with much of the Middle East and North Africa (average Aridity Index=0.05, average tree cover=7%) region and parts of the sub-Saharan Africa region (average Aridity Index=0.5, average tree cover=17%) being arid or semi-arid and having generally low tree cover. Conversely, the Europe region tends to have higher Aridity Index and higher tree cover (average Aridity Index =0.7, average tree cover=25%). Figure 4 demonstrates the difference in distribution of percent coverage across the climate zones which our different study regions fall within.

**Figure 4.**
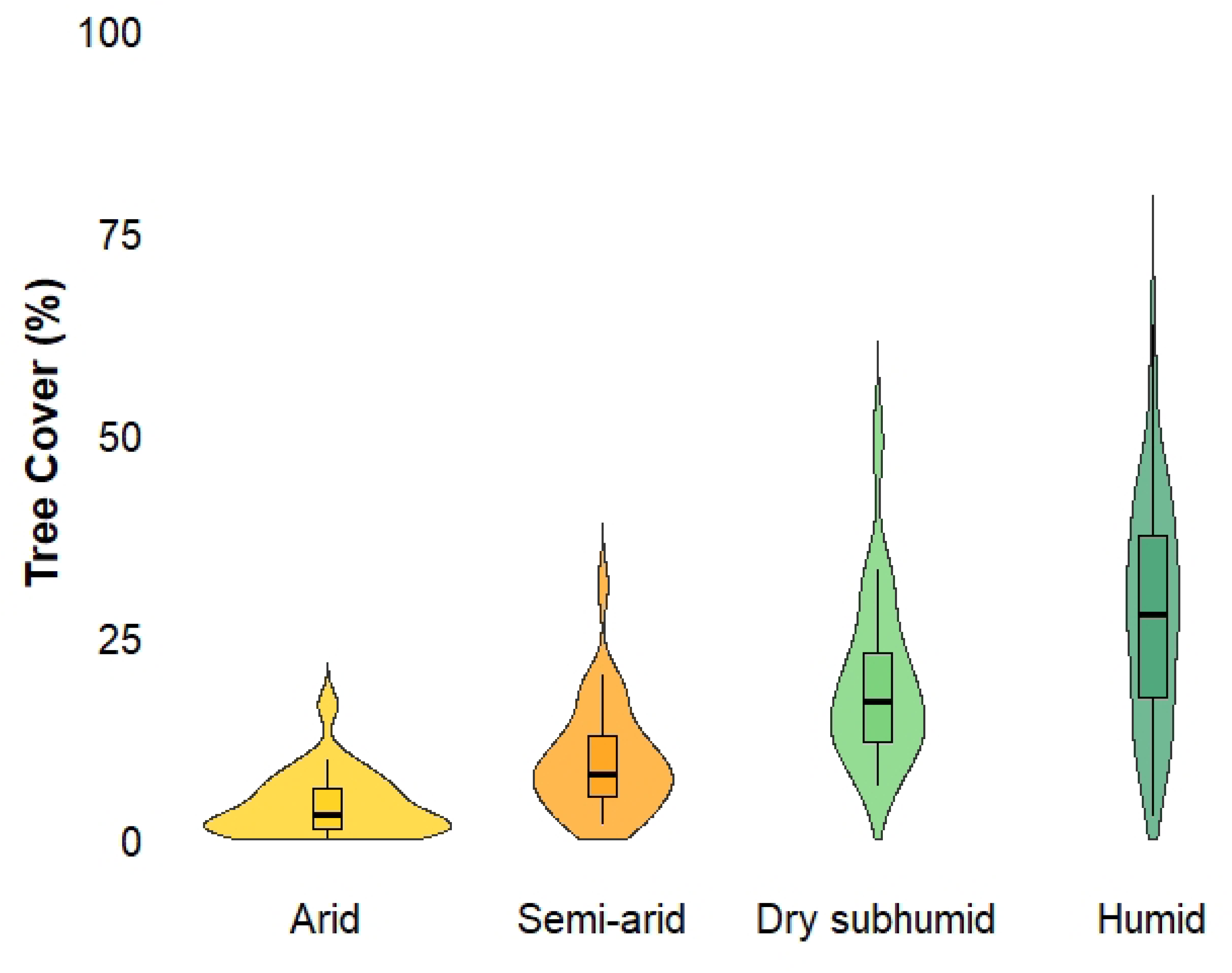
Violin plot of the distribution of tree cover (2021) percentage across four different climate zones: Arid, Semi-arid, Dry Sub-humid, and Humid. Inside the violin plots, there are box plots showing the median and the distribution of the variance.

Other measures of urban nature access considered in this study are only weakly correlated with the aridity index. The average distance to green patches (which can include desert or shrubland land cover types) is not significantly correlated with the aridity index (R= –0.14, p = 0.09). The average distance to OSM parks (which can have varying degrees of vegetation cover) is weakly negatively correlated with the aridity index (R= –0.20, p=0.02).

The population density of an urban area, one simple measure of urban form, in part determines the level of nature access calculated. For instance, population density is negatively correlated (R= –0.39, P < 0.001) with natural cover (Figure 5). In general, higher density FUAs have lower natural cover, presumably because there is more developed cover and thus less natural land cover [19]. However, population density is not significantly related to the distance to OSM parks (R= 0.13 p= 0.12), the distance to green patches (R= 0.06, p= 0.48) or the distance to blue patches (R= –0.09, p= 0.33).

**Figure 5.**
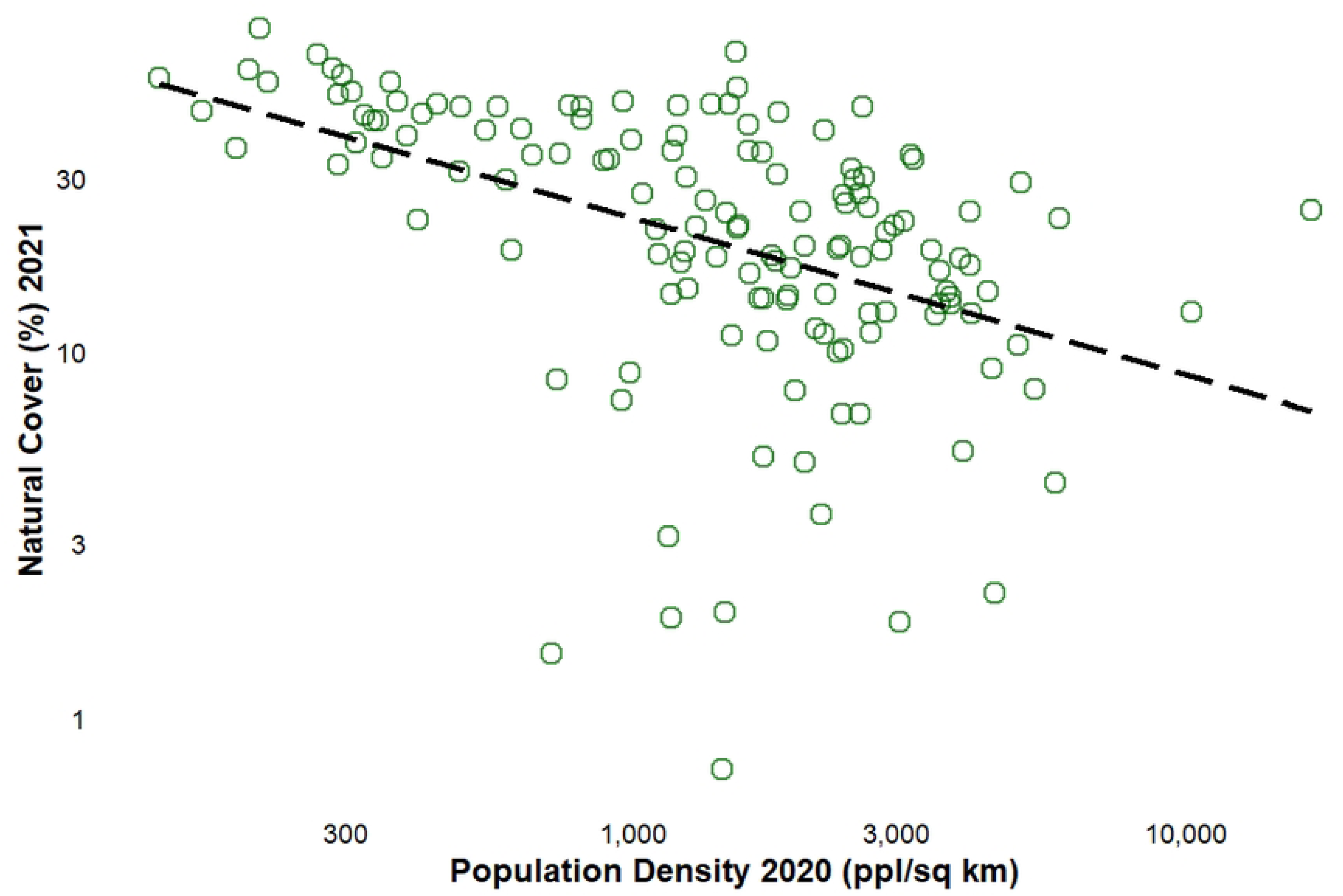
Relationship between population density and weighted natural cover in 2020. Each point represents a Functional Urban Area, with population density (people per square kilometer) on the x-axis and weighted natural cover (%) on the y-axis. Both axes are log-transformed. A linear regression trendline is shown.

Finally, the economic context of urban areas also affects urban nature access. For instance, there is a general negative correlation between Gross Domestic Product (GDP) per capita and the weighted distance to OSM parks (Figure 6). Higher GDP per capita is associated with shorter distances to parks, implying better access to green spaces in wealthier regions. Generally, countries with higher GDP per capita have lower average population density in their FUAs (Figure 6).

**Figure 6.**
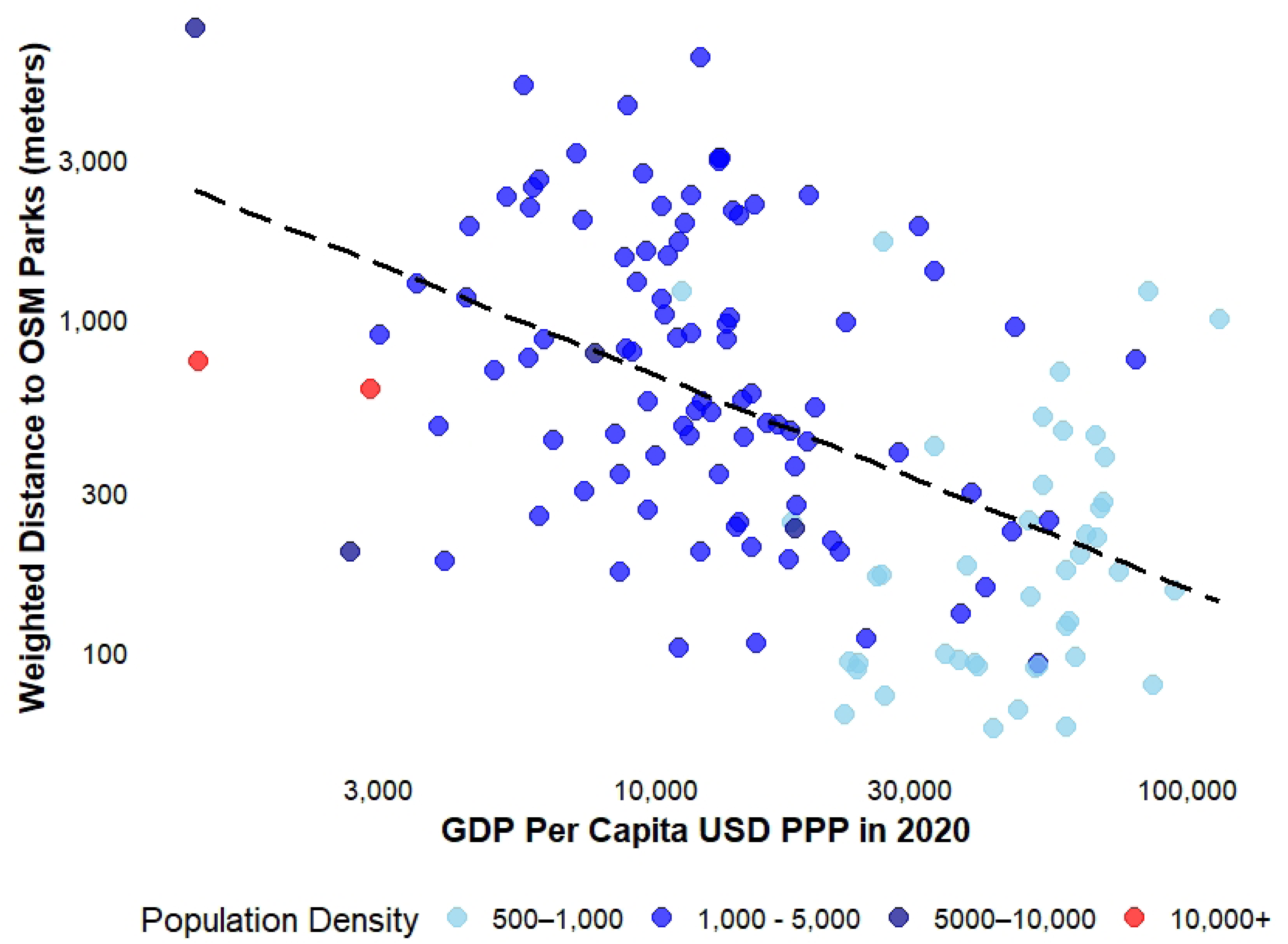
The relationship between GDP per capita and the weighted distance to OSM parks. Note the logarithmic scale on both the X and Y axes. Each Functional Urban Area is represented by circle, with the color of the circle indicating population density (ranging from 500 to 16,000 people per square kilometer).

## Discussion

### Urban access to nature fails to meet benchmarks

After looking at data from a statistically representative sample of global cities, our results show that globally the average urbanite’s exposure and access to nature fails to meet common benchmarks proposed for safeguarding human health and well-being. About 1 in 4 urban dwellers (23% ± 8%) do not live within 1km of an OSM park, some 1.1 billion people across the 4.6 billion people in urban areas worldwide [57]. About 1 in 3 urban dwellers, some 1.6 billion people, do not live within 1km of a green patch (36% ± 9%). The situation is even worse for other metrics, with 3 in 4 urban dwellers (76% ± 7%), some 3.5 billion people, not meeting the 30% threshold for tree cover. Our results are broadly similar to those of Bertassello et al. [32], that examined European cities against the 3/30/300 rule, and found that 21% of Europeans fail to meet any of the three criteria. A substantial scientific literature now shows that exposure and access to nature has a variety of health and well-being benefits [2, 4]. Our results in this paper show that literally billions of urban dwellers have worse health and well-being outcomes because of living in neighborhoods with insufficient nature.

### Declining access over time

Over time, most urban residents have less access to urban nature, at least in terms of access to green patches. With the ESA-CCI dataset, there is a general trend toward a decrease in access to urban nature, at least as measured by large (300m minimum) patches of natural land cover. Between 1992 and 2020, the average distance to a green patch increased for every region. Note that we did find a general trend toward declining distance to blue patches, but more research is needed to verify if this is a real trend, as it seems to be driven by classification differences in the ESA-CCI dataset for specific water features. Regions that had the greatest increase in population density, such as the South Asia region, had the greatest increase in distance (i.e., a decline in nature access) to a green patch. This makes sense, as most FUAs are growing in population, and so their developed area is increasing, which tends to decrease the natural share in the FUA as natural land cover is converted to developed uses. Seen in this light, Target 12 of the Global Biodiversity Framework is then a promise to overcome a global trend, toward decreased nature access over time, at least as measured with this metric.

Overall, our results suggest that it will be difficult to assess urban nature access consistently globally over time, simply because there are few data sources relating to land cover or parks that go back decades and have complete global coverage. In this study, we used the ESA-CCI global dataset, which would be one example of a dataset that allows consistent assessment of urban access over time. There are other land cover datasets that might also be used to monitor over decadal time series, such as land cover classified from Thematic Mapper imagery, which is available from the 1980s onward. We did not use some of the land cover series derived from Thematic Mapper imagery in this project because preliminary inspection suggested that had more temporal inconsistencies in classification than the 300m ESA-CCI data, which also has a greater number of classes that allow for a more nuanced definition of what is natural land cover and what is not. However, there are recent time series of classified Thematic Mapper imagery which may be more temporally consistent [e.g., 58].

It should be noted, however, that other lines of research suggest that urban greenness, as measured through NDVI or other vegetation indices, seems to be increasing in many FUAs, particularly in the Global North [40]. Thus, what may be happening globally is that the average FUA is losing large patches of green space but that many FUAs in the Global North are increasing in urban tree canopy cover and in vegetation associated and intermingled with urban areas (e.g., gardens, small parks, street trees).

### The metric used impacts results

We find the nature access metric used changes the results significantly. This is especially true when looking at the absolute value of a metric, or when comparing that absolute value to an acceptable threshold. For instance, when using parks identified in OSM, there are 23% of all urbanites that are more than 1km from a park, while if land cover data is used to define green patches the statistic is 36%. The OSM dataset is a more complete representation of recreational access than the green patches as defined from land cover, but the land cover data is temporal, allowing estimation of trends in nature access over time, which is impossible with the OSM dataset. To some extent, since many nature access metrics are correlated, even if the absolute value of nature access may vary widely depending on the metric, the rank order of FUAs and regions is similar regardless of the nature access metric used. For instance, regardless of the metric used, the Middle East and North Africa region has low values of nature access compared with other regions. Overall, our results suggest that if policymakers want to assess global progress toward Target 12 in a consistent way, the metric that is selected will substantially affect any quantitative result of how countries/regions are doing in achieving a fixed target, but that the regional patterns in nature access in a comparative sense may be fairly consistent regardless of the metric used.

### Context matters

We find that contextual factors are significantly correlated with the value of nature access metrics and suggest that FUA-specific targets for improvement in urban nature access need to be set after considering contextual factors. For instance, the aridity index is strongly correlated with tree cover in our sample of 140 FUAs, consistent with several earlier studies [34, 59]. This should not be surprising, since trees will only grow without supplemental watering in suitably moist climates, and the same broad climatic gradients that explain the distribution of forested biomes also correlate with patterns of urban forest cover in cities. To a lesser extent, the aridity index is correlated with natural share and distance to green patches. This may be because of the greater forest cover pixels in more mesic FUAs, as forest cover is part of natural share and how we defined green patches. Aridity also correlates with level of economic development (less developed economies are more arid) and with population density (denser FUAs are less green), and these other variables may also be playing a causal role in determining tree cover as well as aridity. If tree cover or related metrics (e.g., NDVI or other measures of greenness from satellite imager) are to be useful for assessing progress to Target 12 of the GBF, then data on tree cover needs to be assessed with this context in mind.

Similarly, we find that less developed economies have cities that tend to have less urban nature access, as measured by distance to OSM park. This is consistent with the findings of Wei et al. [37], who found that urban green space area was greater in developed, wealthier regions like Europe and North America. If distance to parks were to be used to assess progress toward Target 12 of the GBF, then data on distance to park would need to be assessed with this context in mind. It should be emphasized also that while level of economic development is a correlated of distance to OSM park, this correlation is not destiny. There are examples of FUAs in lower-income economies that still have relatively high access to parks, and much can be learned by these “brightspots” [19].

## Conclusions

Many cities globally fail to meet common benchmarks proposed for urban nature access that would be sufficient for safeguarding human health and well-being. Moreover, using one metric, distance to a green space, urban nature access seems to be declining over time. These discouraging trends emphasize why Target 12 and its commitment by global policymakers to increase urban nature access are so important. Our results suggest that it is worthwhile to pick an urban access metric(s) for Target 12 that can be easily assessed globally for many FUAs, over time, such as access to green patches defined from satellite imagery. We would argue that being able to assess progress over time for many cities is worth the loss of precision from using less sophisticated measures of nature access. Our results also suggest that any absolute targets for urban nature access need to be set so that they are considering relevant contextual variables, such as climate, population density, and per-capita GDP. Particularly using nature access metrics derived from land cover, it is now becoming easy with cloud computing and GIS to assess progress for thousands of FUAs at once, allowing the assessment of progress for each and setting targets as some reasonable percentage increase in nature access over the historically defined baselines. Such global, comprehensive assessment of progress toward Target 12 would be valuable, as a supplement to the more detailed data submitted by some cities and countries.

### Contributions

**Robert I. McDonald:** Conceptualization; Methodology; Formal Analysis; Supervision; Writing – original draft. **Mariami Marsagishvili:** Conceptualization; Methodology; Formal Analysis; Writing-review and editing. **Lindsi Seegmiller:** Methodology; Writing-review and editing. **Peter Olsson:** Methodology; Writing-review and editing.

## Data Availability

The datasets created during this research are available on Data Dryad at the following DOI: [Link to Data Dryad will be publicly available after manuscript acceptance. Please let the corresponding authors know if access to this information is necessary during manuscript review, we are happy to provide it.]

